# bigSCale: An Analytical Framework for Big-Scale Single-Cell Data

**DOI:** 10.1101/197244

**Authors:** Giovanni Iacono, Elisabetta Mereu, Amy Guillaumet-Adkins, Roser Corominas, Ivon Cuscó, Gustavo Rodríguez-Esteban, Marta Gut, Luis Alberto Pérez-Jurado, Ivo Gut, Holger Heyn

## Abstract

Single-cell RNA sequencing significantly deepened our insights into complex tissues and latest techniques are capable processing ten-thousands of cells simultaneously. With *big*S*Cale*, we provide an analytical framework being scalable to analyze millions of cells, addressing challenges of future large datasets. Unlike previous methods, *bigSCale* does not constrain data to fit an *a priori*-defined distribution and instead uses an accurate numerical model of noise. We evaluated the performance of *bigSCale* using a biological model of aberrant gene expression in patient derived neuronal progenitor cells and simulated datasets, which underlined its speed and accuracy in differential expression analysis. We further applied *bigSCale* to analyze 1.3 million cells from the mouse developing forebrain. Herein, we identified rare populations, such as *Reelin* positive Cajal-Retzius neurons, for which we determined a previously not recognized heterogeneity associated to distinct differentiation stages, spatial organization and cellular function. Together, *bigSCale* presents a perfect solution to address future challenges of large single-cell datasets.

**Extended Abstract:** Single-cell RNA sequencing (scRNAseq) significantly deepened our insights into complex tissues by providing high-resolution phenotypes for individual cells. Recent microfluidic-based methods are scalable to ten-thousands of cells, enabling an unbiased sampling and comprehensive characterization without prior knowledge. Increasing cell numbers, however, generates extremely big datasets, which extends processing time and challenges computing resources. Current scRNAseq analysis tools are not designed to analyze datasets larger than from thousands of cells and often lack sensitivity and specificity to identify marker genes for cell populations or experimental conditions. With *big*S*Cale*, we provide an analytical framework for the sensitive detection of population markers and differentially expressed genes, being scalable to analyze millions of single cells. Unlike other methods that use simple or mixture probabilistic models with negative binomial, gamma or Poisson distributions to handle the noise and sparsity of scRNAseq data, *bigSCale* does not constrain the data to fit an *a priori*-defined distribution. Instead, *bigSCale* uses large sample sizes to estimate a highly accurate and comprehensive numerical model of noise and gene expression. The framework further includes modules for differential expression (DE) analysis, cell clustering and population marker identification. Moreover, a directed convolution strategy allows processing of extremely large data sets, while preserving the transcript information from individual cells.

We evaluate the performance of *bigSCale* using a biological model for reduced or elevated gene expression levels. Specifically, we perform scRNAseq of 1,920 patient derived neuronal progenitor cells from Williams-Beuren and 7q11.23 microduplication syndrome patients, harboring a deletion or duplication of 7q11.23, respectively. The affected region contains 28 genes whose transcriptional levels vary in line with their allele frequency. *BigSCale* detects expression changes with respect to cells from a healthy donor and outperforms other methods for single-cell DE analysis in sensitivity. Simulated data sets, underline the performance of *bigSCale* in DE analysis as it is faster and more sensitive and specific than other methods. The probabilistic model of cell-distances within *bigSCale* is further suitable for unsupervised clustering and the identification of cell types and subpopulations. Using *bigSCale*, we identify all major cell types of the somatosensory cortex and hippocampus analyzing 3,005 cells from adult mouse brains. Remarkably, we increase the number of cell population specific marker genes 4-6-fold compared to the original analysis and, moreover, define markers of higher order cell types. These include CD90 (*Thy1*), a neuronal surface receptor, potentially suitable for isolating intact neurons from complex brain samples.

To test its applicability for large data sets, we apply *bigSCale* on scRNAseq data from 1.3 million cells derived from the pallium of the mouse developing forebrain (E18, 10x Genomics). Our directed down-sampling strategy accumulates transcript counts from cells with similar transcriptional profiles into index cell transcriptomes, thereby defining cellular clusters with improved resolution. Accordingly, index cell clusters provide a rich resource of marker genes for the main brain cell types and less frequent subpopulations. Our analysis of rare populations includes poorly characterized developmental cell types, such as neuron progenitors from the subventricular zone and neocortical *Reelin* positive neurons known as Cajal-Retzius (CR) cells. The latter represent a transient population which regulates the laminar formation of the developing neocortex and whose malfunctioning causes major neurodevelopmental disorders like autism or schizophrenia. Most importantly, index cell cluster can be deconvoluted to individual cell level for targeted analysis of populations of interest. Through decomposition of *Reelin* positive neurons, we determined a previously not recognized heterogeneity among CR cells, which we could associate to distinct differentiation stages as well as spatial and functional differences in the developing mouse brain. Specifically, subtypes of CR cells identified by *bigSCale* express different compositions of NMDA, AMPA and glycine receptor subunits, pointing to subpopulations with distinct membrane properties. Furthermore, we found *Cxcl12*, a chemokine secreted by the meninges and regulating the tangential migration of CR cells, to be also expressed in CR cells located in the marginal zone of the neocortex, indicating a self-regulated migration capacity.

Together, *bigSCale* presents a perfect solution for the processing and analysis of scRNAseq data from millions of single cells. Its speed and sensitivity makes it suitable to the address future challenges of large single-cell data sets.

## Introduction

Single-cell RNA sequencing (scRNAseq) is at the forefront of techniques to chart molecular properties of individual cells. Recent microfluidic-based methods are scalable to ten-thousands of cells, enabling an unbiased sampling and in-depth characterization without prior knowledge^1–3^. Consequently, studies are less confined by the number of cells and aim to produce comprehensive cellular atlases of entire tissues, organs and organisms^4^. Increasing cell numbers, however, generate extremely large datasets, which extend processing time and challenge computing resources. Current scRNAseq analysis tools are not designed to analyze datasets larger than thousands of cells and often lack sensitivity and specificity to identify marker genes for cell populations or experimental conditions.

To address the challenges of large scRNAseq datasets, we developed *big*S*Cale*, an analytical framework for the sensitive detection of population markers and differentially expressed genes, being scalable to analyze millions of single cells. Unlike other methods that use simple or mixture probabilistic models with predefined distributions to handle the noise and sparsity of scRNAseq data^5–8^, *bigSCale* does not assume an *a priori*-defined distribution. Instead, *bigSCale* uses large sample sizes to estimate a highly accurate and comprehensive numerical model of noise. The framework further includes modules for differential expression (DE) analysis, cell clustering and population marker identification. Moreover, a directed convolution strategy allows the processing of extremely large datasets, while preserving the transcript information from individual cells.

We evaluate the performance of *bigSCale* using a defined biological model for reduced or elevated gene expression levels by performing scRNAseq of neuronal progenitors derived from induced pluripotent stem cells of Williams-Beuren^9^ and 7q11.23 microduplication^10^ syndrome patients. Simulated datasets of different size and sparsity were utilized to underline the accuracy and speed of *bigSCale* in DE analysis. To demonstrate its suitability for unsupervised clustering and population marker identification using its probabilistic model of cell-distances, we applied *bigSCale* to cluster cell types of the somatosensory cortex and hippocampus from adult mouse brains^11^. Lastly, the *bigSCale* framework was applied to convolute and characterize 1.3 million cells derived from the developing mouse forebrain, detecting profound heterogeneity in rare neuronal subpopulations. We believe *bigSCale* presents a perfect solution for the processing and analysis of scRNAseq data from millions of single cells. Its speed and sensitivity make it suitable to address future challenges of large single-cell datasets.

## Results

### The *bigSCale* framework

Datasets from scRNAseq display sparse and noisy gene expression values, among other sources due to drop-out events, amplification biases, and variable sequencing depth. The *bigSCale* framework builds a probabilistic model to define phenotypic distance between pairs of cells that considers all sources of variability. Compared to other methods that assume negative binomial, gamma or Poisson distributions in simple or mixture probabilistic models, *bigSCale* estimates a highly accurate and comprehensive numerical model of noise. The model allows to quantify distances between cells, which provide the basis for differential expression analysis and cell clustering (**Fig. 1**, **Methods**).

**Fig.1.**
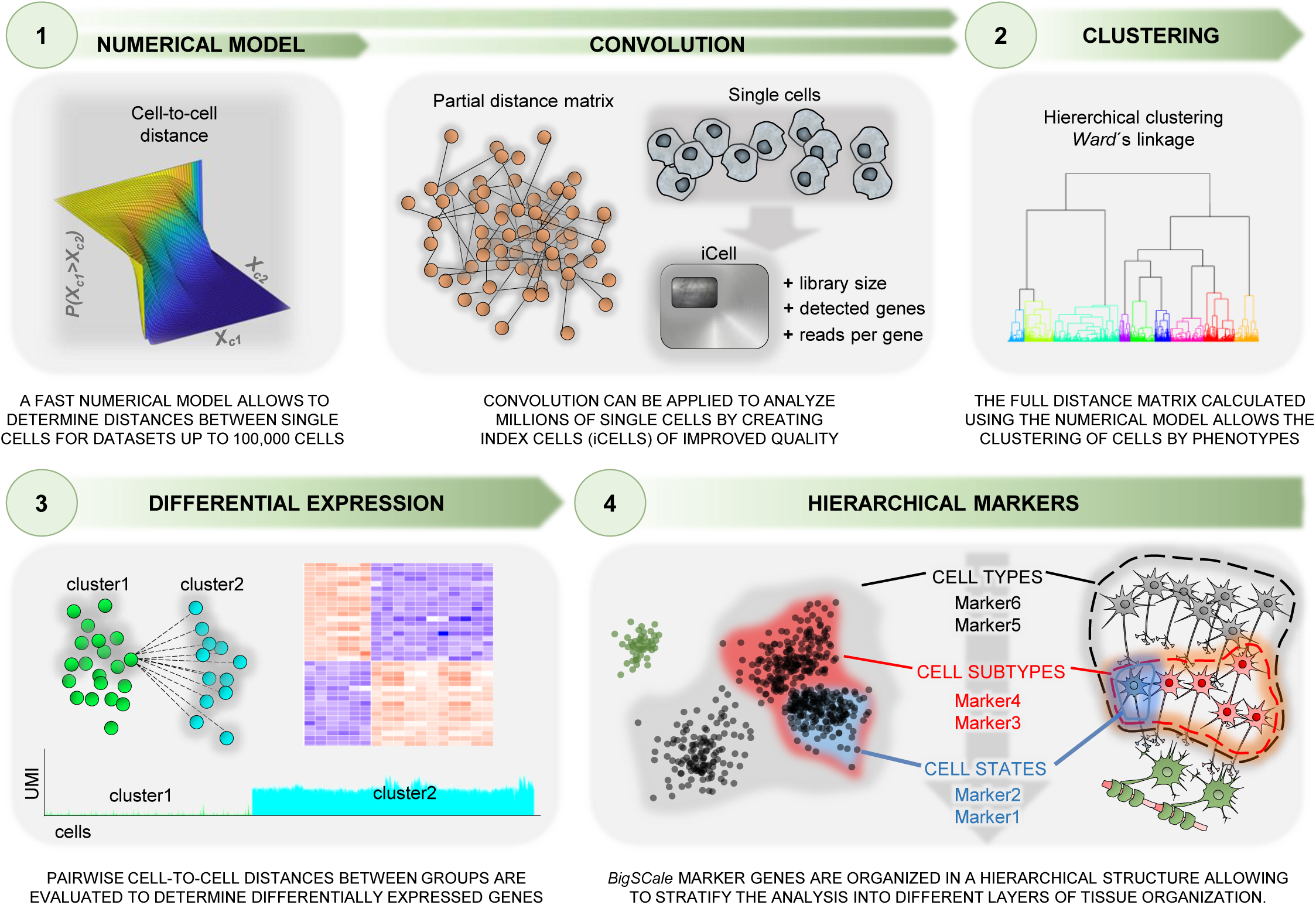
Schematic representation of the *bigSCale* framework for analyzing millions of single-cell transcriptomes. The analytical framework includes a numerical model step to determine distances between single cells and modules for differential expression analysis, cell clustering and population marker identification. An optional convolution strategy allows the processing of extremely large datasets (preserving the transcript information from individual cells).

**(1)** To generate the model, cells featuring highly similar transcriptomes are grouped together. Next, the expression variation within groups is used as an estimator of noise. Unlike previous methods, *bigSCale* models differences in expression levels rather than expression levels themselves. Therefore, a p-value is assigned to each gene, representing the likelihood of a change of expression from one cell to another. Prior to model computation, a module for batch effect removal can be applied.

**(2)** For differential expression *bigSCale* assigns a p-value to each gene, representing the likelihood of an expression change between two groups of cells. To this end, all pairwise cell comparisons between two groups are performed. Genes repeatedly differing in expression between cells cumulate higher scores, which are next adjusted and normalized into p-values.

**(3)** Cellular clustering is achieved by computing all pairwise cell distances to generate a distance matrix and to assign cells into groups (via *Ward’s* linkage). Specifically, the distance matrix is computed over a set of overdispersed genes, namely genes presenting a high degree of variation across the dataset. To improve the feature quality, skewed, isolated and perfectly correlating genes are discarded. The latter are prone to generate artificial transcript clusters and consist of genes with a common 3’-end, being indistinguishable by digital counting scRNAseq methods.

**(4)** Following the identification of cell clusters, *bigSCale* conducts an iterative DE analysis between populations of cells for the sensitive detection of markers, defined by genes unevenly expressed across populations. Notably, most current tools lack the option to model multifaceted phenotype structures with overlying molecular signatures of cells. Conversely, *bigSCale* allows to disclose multiple alternative phenotypes of a given cell by ordering markers in a hierarchical structure, in which increasing layers of phenotypic complexity (from cell types to subtypes or states) are represented by markers at increasing hierarchical levels.

**(5)** While *bigSCale’s* intrinsic speed allows the direct analysis of datasets up to hundred thousand cells, adjustments are needed to handle millions of cells. For these scenarios, the cell numbers are scaled down by pooling (convoluting) information from cells with analogous transcriptional profiles into indexed cell (iCell) profiles. Here, iCells are defined by adding transcript counts from pools of similar single cells, significantly increasing molecule and gene counts and overall improving the expression profile quality. Accordingly, iCells allow to discriminate subpopulations with higher precision and sensitivity. Most importantly, iCells preserve the transcript information from individual cells and can be deconvoluted for targeted analysis of populations of interest.

### Identification of differentially expressed genes

We evaluate the performance of *bigSCale* using a biological model for reduced and elevated gene expression levels. Specifically, we performed scRNAseq of 1,920 neuronal progenitor cells (NPC) derived from induced pluripotent stem (iPS) cells of two patients with Williams-Beuren (WB) and two with 7q11.23 microduplication (Dup7) syndrome. Both are multisystemic disorders caused by a heterozygous deletion or duplication, respectively, of 1.5-1.8Mb at the chromosome band 7q11.23. This region is flanked by segmental duplications with high sequence identity that can mediate nonhomologous recombination with the consequent loss or gain of 26-30 contiguous genes, whose transcriptional levels vary in line with their allele dosage^9,10^. To benchmark *bigSCale* against other common single-cell DE tools, NPC from four syndromic patients (WB1/2: n=742 and Dup7.1/2: n=735) were compared to NPC derived from a healthy donor (WT: n=369 cells). The sensitivity of each algorithm was evaluated by counting the number of genes detected to be significantly down- or upregulated in patients against the control. To achieve the same level of specificity amongst tools, the top 1500, 2000 and 2500 deregulated genes were used in each comparison.

For the WB1 sample harboring a deleted allele, *bigSCale* presented the highest sensitivity by detecting 12 down-regulated genes, followed by *scde*^6^, *MAST*^7^, *seurat*^5^ and *scDD*^8^ (**Fig. 2a**). Notably, *bigSCale* finds the same genes as the other best performing tools, plus additional events (**Fig. 2b**). Interestingly, the poorest performing tool *scDD* is also the most divergent one, displaying reduced overlaps with the other four methods (**Fig. 2a,b**). Consistently, *bigSCale* displayed the highest sensitivity also in the remaining three comparisons (**Supplementary Fig. 1a-c**), with an overall average of 11.5 detected down-regulated genes in WB patients and 9 up-regulated genes in Dup7 patients (**Fig. 2c**). Moreover, *bigSCale* proved to be the most sensitive method at all tested specificity levels, with an average of 8.75 (top 2000) and 6.75 (top 1500) detected DE genes (**Supplementary Fig. 1d**). These results indicate that *bigSCale* outperforms other methods for single-cell DE analysis in sensitivity when using biological data.

**Fig.2.**
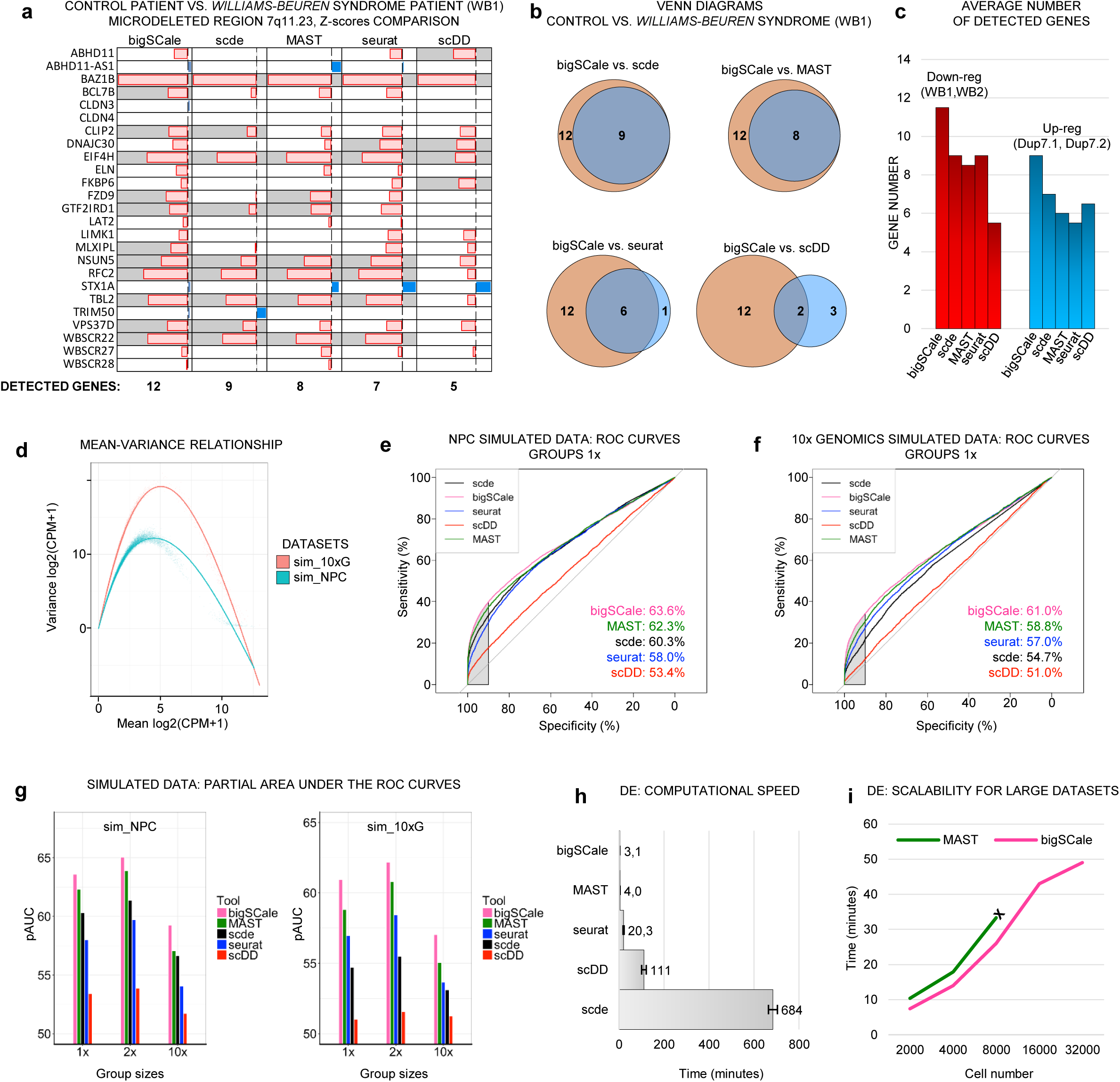
Benchmarking of sensitivity, specificity and speed of *bigSCale*, *scde*, *seurat*, *MAST* and *scDD*. (**a**) Differential expression analysis in iPS cell-derived neuronal progenitor cells (NPC) from healthy and Williams-Beuren (WB) syndrome donors (WT vs. WB1). For the genes located in the deleted region the p-values of each tool are shown in Z-score scale, (blue: down-regulated; red: up-regulated). Genes correctly detected as down-regulated are highlighted (grey). Total numbers of correctly assigned genes are indicated (below). (**b**) Venn diagrams for WT vs. WB1 comparing the identity of correctly assigned genes (orange: *bigSCale*; blue: others) (**c**) Average number of detected down-(blue) and up-regulated (red) genes in the two WB and Dup7 patients, respectively, compared to healthy donor. (**d**) Comparison of the mean-variance relationship in the two simulated datasets (sim_NPC and sim_10x). (**e,f**) Partial AUCs of ROC curves computed across the tools in the two simulated datasets (sim_NPC, **e**; sim_10x, **f**) with group sizes having proportions 1:1 (1x). The sensitivity at high level of specificity (>90%) is highlighted (grey area). (**g**) Barplots of partial AUC across tools for all tested proportions (1x, 2x, 10x) in DE analyses of simulated datasets (sim_NPC and sim_10x). **h**) Average required time for computing DE in the NPC cell model (average 739 total cells per comparison, 4 comparisons, tools run on 1 CPU-core) **i**) Scalability of *bigSCale* and *MAST* with large datasets. *MAST* could not be tested beyond 8,000 cells due to excessive RAM requirements (>16Gb).

To further test the performances in determining DE genes, we benchmarked *bigSCale* against the previous tools using simulated datasets. For data simulation, we used *Splatter*^12^, which allows to generate and control true positive DE genes. Simulations have been performed estimating parameters from two datasets representing different characteristics of large-scale experiments, namely our NPC dataset (sim_NPC) and a droplet-based experiment consisting of ~2,500 cells sequenced to low coverage (10x Genomics, sim_10xG; **Methods**). The two datasets widely differed in the number of detected genes per cell, sparsity and heterogeneity (**Fig. 2d** and **Supplementary Fig. 2a**). In both simulations, we recreated distributions of gene expression levels and library properties highly similar to the original datasets and preserved the original number of cells and genes. Six cell types of different proportions were simulated in each dataset, allowing to test DE between groups of proportions 1:1 (1x), 1:2 (2x) and 1:10 (10x). Each tool has been applied to the complete dataset at the model-building step, prior to test DE between groups of cells.

The ability to correctly classify true DE genes against non-DE genes was evaluated calculating the area under the curve (AUC) of a receiver operating characteristic (ROC) curve, ranking genes in their order of significance as determined by the tools. To test the capacity of controlling false positives events, we focused on the partial AUC with high specificity being >90%. All tools performed better in the simulated NPC dataset and the order of tools was consistent across all group sizes (**Fig. 2e,f** and **Supplementary Fig. 2b-e**). Remarkably, *bigSCale* outperformed the other tools, reaching the highest levels of sensitivity and specificity in all tested conditions (**Fig. 2g**). The *MAST* performance was the closest to *bigSCale*, with the gap being more evident in more distinct proportional contexts (10x, **Supplementary Fig. 2c,e**). Notably, while the design of *MAST* is restricted to DE analysis, *bigSCale* provides a comprehensive framework for single-cell analysis.

In the view of increasing datasets sizes, we further evaluated *bigSCale*’s speed in DE analysis. In the biological model (NPC), *bigSCale* proved to be the fastest tool (3.1 min) in performing DE, followed by *MAST* (4.0 min) (**Fig. 2h**). The slowest tool was *scde* (684 min), as reported in previously studies^13,14^. We next compared the scalability of *bigSCale* to *MAST* with respect to samples sizes. To this end, we created a simulated matrix of 40,000 genes and 32,000 cells and performed DE analyses between pairs of groups with sizes ranging from 2,000 to 32,000 cells. *BigSCale* was faster for all conditions (**Fig. 2i**). Moreover, *bigSCale* could process datasets larger than 8,000 cells, whereas *MAST* was limited by the RAM requirements, denoting a broader perspective of applicability for *bigSCale*.

### Cellular clustering and population marker identification

To evaluate the ability of *bigSCale* to identify cell types and subpopulations in complex tissues, we analyzed 3,005 cells of the somatosensory cortex and hippocampus dissected from the adult mouse brains^11^. Consistently with previous analyses^11,15^, *bigSCale* was able to segregate all major brain cell types, namely somatosensory pyramidal neurons, different types of CA1/2 pyramidal neurons, interneurons, astrocytes, oligodendrocytes and vascular cells (**Fig. 3a**). Remarkably, we increased the number of marker genes specific for cell types 4/5-fold compared to the original analysis using *BackSPIN* and, moreover, defined markers of higher order cell types (**Fig. 3a**). Specifically, *bigSCale* determined 9,258 marker genes for cellular types, including 7,167 previously unidentified markers (**Supplementary Table 1**). The expression patterns of the novel markers were highly specific to the respective populations of cells, as shown for astrocytes (**Fig. 3b**), oligodendrocytes, vascular cells, neurons and interneurons (**Supplementary Fig. 3a,b**), pointing to a high accuracy of *bigSCale*. In line, external bulk RNAseq signatures supported the novel markers to be highly specific for the respective populations^16,17^ (astrocytes, p<4.9e-62; oligodendrocytes, p=9.9e-18; interneurons, p=9.8e-19; neurons, p=2.3e-34; vascular, p=1.0e-67). Furthermore, the novel markers included established marker for brain subtypes, such as *Atp1a2*^18^, *Slc1a3*^19^, *Mt1*^20^ and *Aqp4*^21^ for astrocytes or *Stmn3*^22^, *Snap25*^23^ for neurons (**Supplementary Fig. 4a-c**).

**Fig.3.**
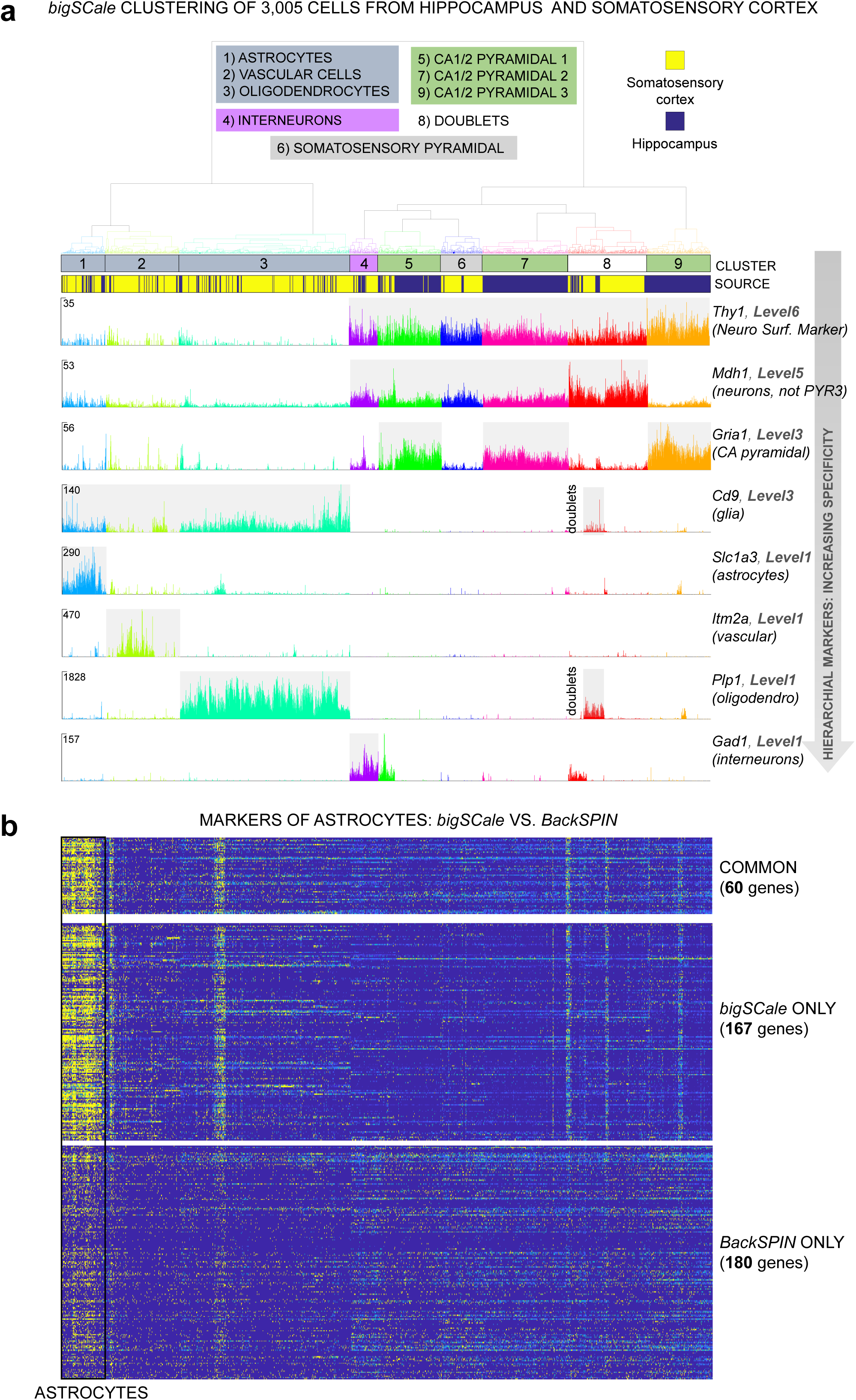
*BigSCale* analysis of scRNAseq data from 3,005 mouse cortical and hippocampal cells^11^. (**a**) Dendrogram and expression plots reporting examples of hierarchical markers. Dendrogram was cut at 20% of its total depth to segregate 9 different clusters of cells, which correspond to the main brain cell types. In the expression plots, UMI counts are shown at single-cell level for markers of different hierarchical marker levels (Online Methods). Marker genes for decreasing marker levels, representing distinct brain cell types are displayed. (**b**) Comparison of *bigSCale* and *BackSPIN*^11^ in the detection of gene markers for astrocytes. *BigSCale* identified 167 additional markers with high specificity for astrocytes (high expression, yellow; low expression, blue). *Vice versa*, markers uniquely identified by *BackSPIN* display a weak specificity and achieved low scoring in *bigSCale*.

Differently to other methods, *bigSCale* marker genes are organized in a hierarchical structure allowing to stratify the analysis into different layers of tissue organization. This enabled the assignment of markers to subpopulations, but also higher order cell types, such as glia cells or neurons (**Supplementary Fig. 3b**). In this regard, current experimental designs fail to reliably separate intact neurons from glia cells, as established markers (e.g. *NeuN*) located in the nuclear membrane and are not suitable for isolating entire neurons. Our analysis identified 1,656 marker genes silenced in glial cells and expressed in neuronal populations (**Supplementary Table 1**), such as the neuronal surface receptor CD90 (*Thy1,* **Fig. 3a**), potentially suitable for isolating intact neurons from complex brain samples.

### Convolution of large datasets into index cells

To analyze very large datasets of millions of cells, *bigSCale* convolutes the original cells into iCells with improved transcriptional profiles after the numerical model has been computed using the entire dataset (**Methods**). To ensure that the convolution strategy does not deteriorate cellular phenotypes and related cell clustering, we evaluated its performance by analyzing 20,000 brain cells (randomly downsampled data set, 10x Genomics). Specifically, we tested the cluster assignment of all cell pairs within the dataset before and after increasing degree of convolution (from 4,587 to 2,101 iCells) and for different cluster numbers (n=2-32). Similarities of classification were defined by the *Rand Index* (*RI*), a metric suitable for comparative cluster assessment^24^, where *RI*=100% implies complete similarity of clusterings. Importantly, we observed a highly similar cluster assignment between original and convoluted datasets with *RI*>80% (**Fig. 4a**). The RI was also stable with increasing cluster numbers or degree of convolution, indicating a robust strategy to reduce cell numbers. In line, visualizing cells in two-dimensional plots (tSNE) confirmed the high similarity of cluster assignment between original and iCells (**Fig. 4b**). Together, the results support the utility of *bigSCale* convolution to reduce dataset sizes without the introduction of artifacts.

**Fig.4.**
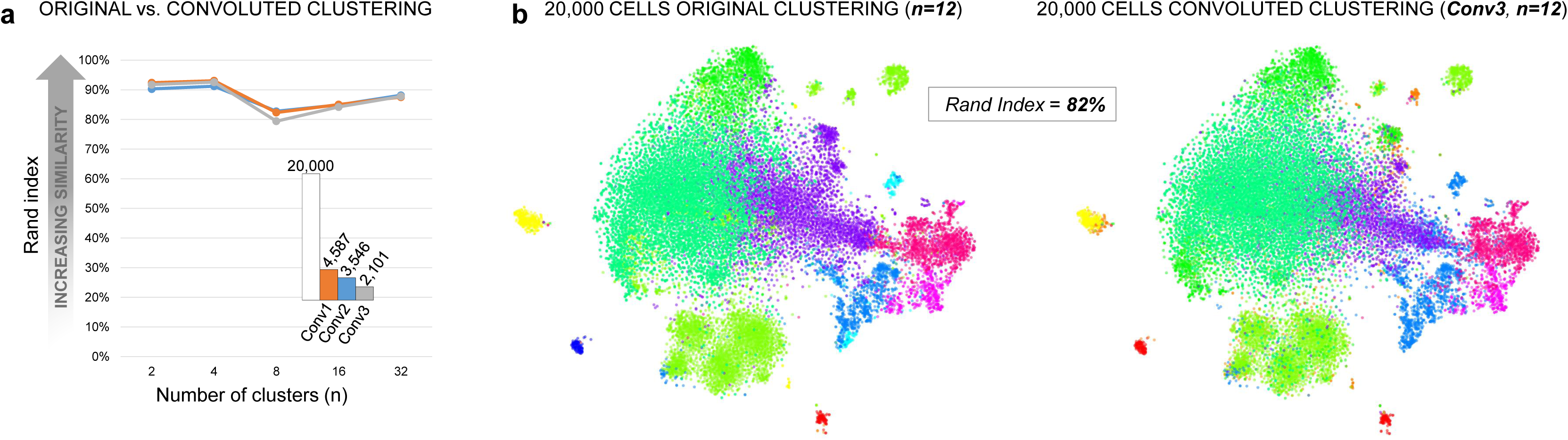
Assessment of the cell convolution strategy in *bigSCale*. (**a**) Comparison of original and convoluted clustering with the Rand-index. Pairwise cell comparisons were performed for three increasing degrees of convolution (Conv1,2,3) into iCells (numbers indicated). Similarity of clustering (Rand-index, y axis) were evaluated at different resolution (n cluster numbers, x axis). Rand-indexes were >80% for all tested combinations, pointing to highly similar cluster assignment for original and iCells. (**b**) tSNE plots comparing original and convoluted clustering. The example displays a comparison with Rand-index = 82% and 12 clusters. The high degree of concordance between experiments is visible through the consistent cluster assignment of cell pairs.

### Analysis of 1,306,127 cells of the developmental pallium

The by far most extensive dataset to date for scRNAseq are 1,306,127 sequenced mouse brain cells from the developmental (E18) dorsal and medial pallium. The data was produced using droplet-based library preparation (10xChromium v2) and is publically available (10x Genomics). Despite being the sole developmental scRNAseq dataset of crucial regions such as cortex, hippocampus and the subventricular zone, its large size yet prevented any detailed analysis. We reasoned that the *bigSCale* analytical framework would be suitable to analyze such large data set and performed an in-depth analysis of cell types and states, including rare and poorly described subpopulations. This analysis serves as proof-of-concept for *bigSCale*’s suitability to process millions of cells from complex tissues in an unbiased manner.

Initially, we applied our convolution strategy to reduce the dataset size 50-fold from 1,306,127 cells to 26,185 iCells. As expected iCells were of improved quality with average library size increasing 50-fold (from 4,890 to 238,500 UMIs) and detected genes per cell increasing 5-fold (from 2,009 to 9,360). In line, average expression level increased from to 2.4 UMIs to 25.5 UMIs. The convolution retained 1,244,298 cells (95.27%), discarding 61,829 cells (4.73%). Clustering of the index cells revealed 16 major cell populations and captured 16,242 differentially expressed markers (**Fig. 5a,b**, **Supplementary Table 2**). We classified the 16 populations in four main cell types: non-neuronal (1-4), neuronal progenitors (5-8), radial glia (9-11) and post-mitotic neurons (12-16). Hierarchical markers allowed to sharply disentangle cell types and subtypes, as well as stages of lineage commitment. Exemplarily, higher order markers, such as *Tubb3* and *Slc1a3* mark the two main cell types: postmitotic neurons of the intermediate/marginal zones and radial glia and progenitors of the ventricular zone, respectively (**Fig. 5a,c**). Similarly, *bigSCale* captured the hallmarks of the main stages of the neuronal lineage^25^, indicated by the expression of *Pax6* (radial glia), *Tbr2* (committed progenitors) and *Tbr1* (differentiated neurons) (**Fig. 5a**). On the other hand, the most significant markers shaping the heterogeneity of post-mitotic neurons are *Stmn2* (silenced in neuroblasts), *Meg3* (interneurons and Cajal-Retzius (CR) neurons), *Nrp1* (glutamatergic neurons), *Tac2* (neuroblasts), *Reln* (CR neurons) and *Gad2* (gabaergic interneurons).

**Fig.5.**
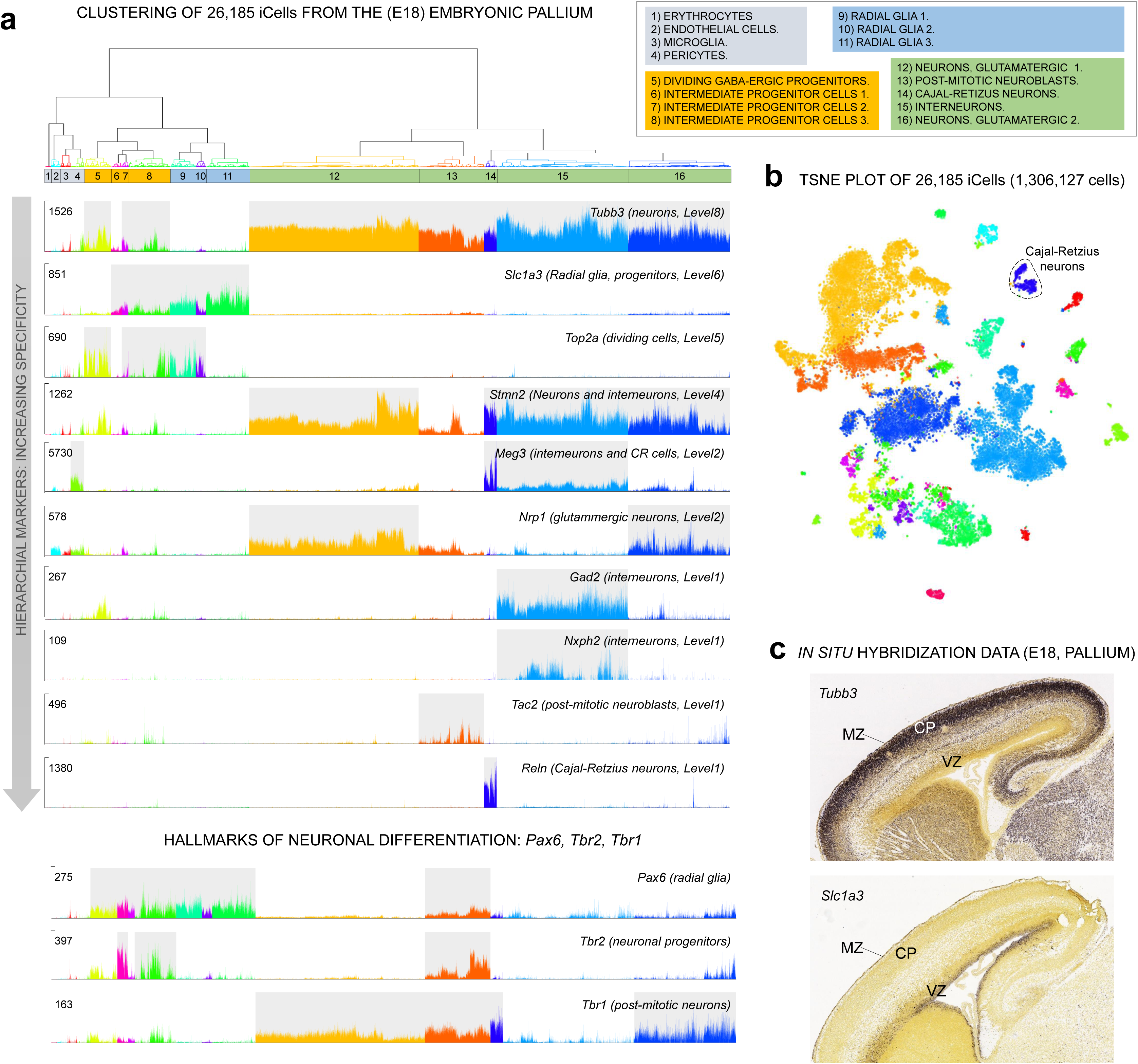
*BigSCale* analysis of 26,185 iCells (convoluted from 1,306,127 single cells) of the embryonic pallium (E18). (**a**) Dendrogram of 16 iCell clusters representing the major cell types (split by color) and subpopulations (cluster 1-16). Single-cell expression plots (UMI counts) present marker genes (decreasing levels of hierarchical markers) for the main subpopulations and specific markers for neuronal differentiation (lower panel). (**b**) tSNE representation of the 16 populations of pallial cells identified by *bigSCale* clustering. (**c**) In-situ hybridization data for *Tubb3* and *Slc1a3*. Post-mitotic neurons (*Tubb3* positive) locate to the outer neocortical layers, including cortical plate (CP) and marginal zone (MR) and radial glia and progenitors (*Scl1a3* positive) are found in the ventricular and sub-ventricular zone (VZ).

As expected, some radial glia (C9, C10) and progenitor populations (C5, C7, C8) represent dividing cells, indicated by *Top2a* expression and other cell cycle genes (**Fig. 5a**, **Supplementary Table 2**). Interestingly, *bigSCale* also identified a population of dividing GABAergic progenitors (C5) characterized, amongst other markers, by simultaneous expression of *Gad2, Pax6* and *Top2a*. Subpatterns of expression within populations of cells further indicate the presence of subtypes of cells, as displayed by the uneven expression of the signaling molecule *Nhpx2* within *Gad2* positive interneurons (**Fig. 5a**). Given the association of *Nhpx2* with an attention-deficit/hyperactivity disorder^26^, *Gad2/Nhpx2* positive cells could represent a previously unknown developmental subtype of interneurons with a roles in behavior and neurocognitive functions.

### Deconvolution for high-resolution subpopulation analysis

While *bigSCale* enabled the convolution of 1.3 million cells to characterize the main cellular types of the developmental pallium with unprecedented detail, the information of single-cell transcriptional profiles was maintained. Consequently, population specific deconvolution allows the in-depth analysis of populations of interest at the resolution of individual cells. We were especially interested in the population of *Reln* positive cells, also known as Cajal-Retzius neurons, a transient type of neurons which regulates the laminar formation of the developing neocortex and whose malfunctioning causes major neurodevelopmental disorders like autism or schizophrenia^27^.

To date a comprehensive phenotypic characterization of the CR cells and its potential subtypes remains elusive, mostly due to their transient nature and to the lack of unambiguous markers. To unravel the diversity of CR cells, we deconvoluted 480 *Reln* positive iCells to 17,543 individual *Reln* positive cells, an unprecedented resource to phenotype this cell type (**Fig. 6a**). *Reln* was expressed uniformly in all deconvoluted cell, confirming the specificity of the convolution strategy (**Fig. 6b**). Furthermore, P73 (Trp73) a well-known marker of neocortical CR cells of later developmental stages (E18) was also uniformly expressed. Expression of P73 indicates that the CR cells were originated from the cortical hem, which is the major source of neocortical CR cells^28^. We determined CR cell specific markers, in addition to *Reln* and *Trp73*, which included *Cacna2d2*, a calcium channel subunit, and *Eya2,* a transcriptional coactivator (**Methods**). Unsupervised clustering revealed eight major subpopulations of CR cells (**Fig. 5a**) and a total of 8,174 differentially expressed markers genes (**Supplementary Table 3**). The clusters also included cell doublets an inevitable artefact of microfluidic-based sample processing, recognizable by cells with simultaneous expression of *Reln* and erythrocytes genes (**Fig. 6a**).

**Fig.6.**
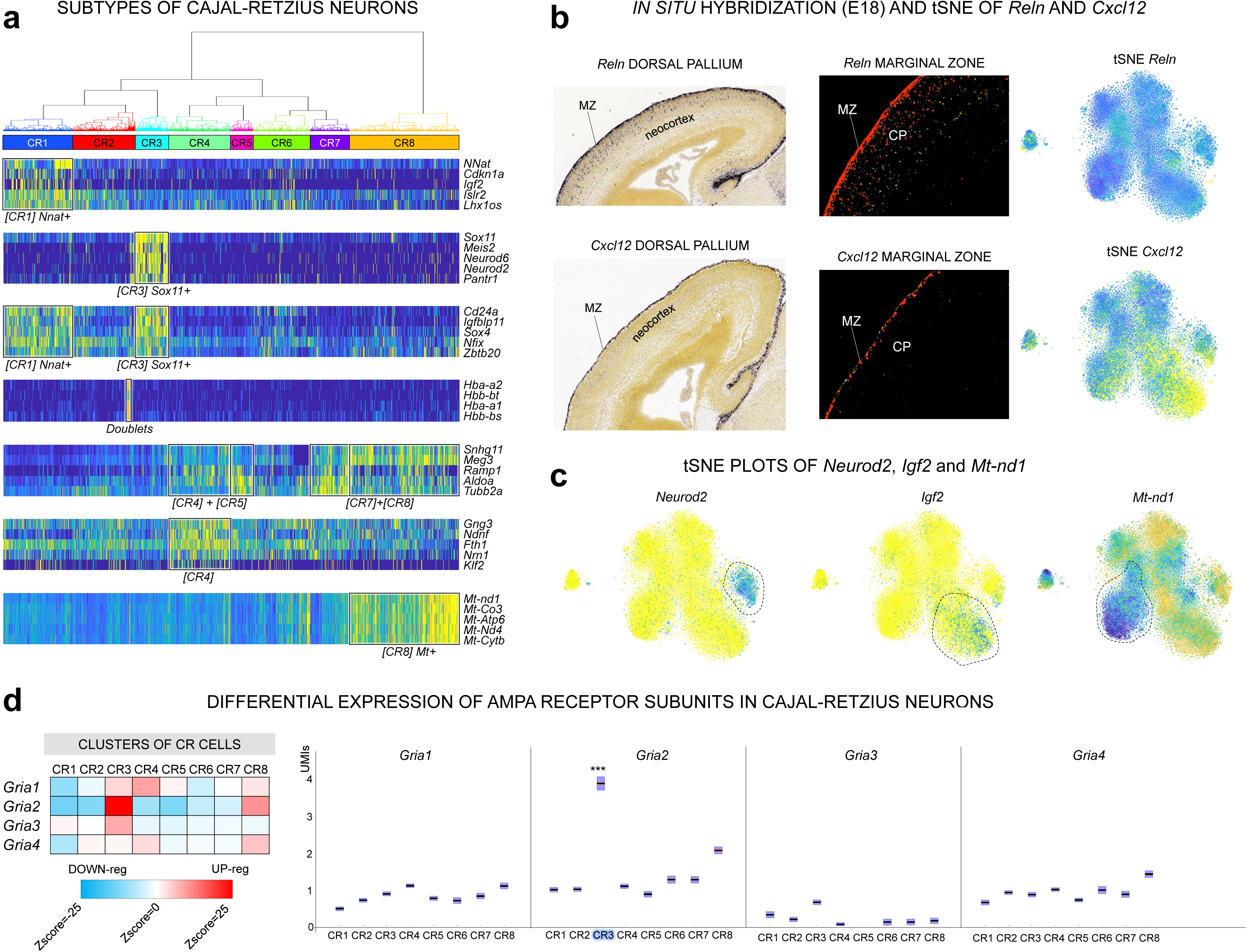
Subtypes of Cajal-Retzius (CR) cells disentangled by *bigSCale*. (**a**) Dendrogram and heatmap of the five top-scoring population markers (CR1-8; high expression, yellow; low expression, blue). (**b**) Comparison of *Reln* (upper panel) and *Cxcl12* (lower panel) expression spatially resolved. *Reln* consistently marks all CR cells, (tSNE, right) located in the Marginal Zone (MZ) and the Cortical Plate (CP) *in situ* immuno-(left) and fluorescence-staining (middle, source: Allen Mouse Brain). *Cxcl12* is expressed in a CR subpopulation (tSNE, right and *in situ* experiments indicate that *Cxcl12* positive cells are exclusively located in the marginal zone. (**c**) tSNE representation of *Neurod2*, *Igf2* and *Mt-nd1* positive subpopulations of CR cells. (**d**) Differential expression of AMPA receptor subunits in CR cells. (left) Heatmap (Z-scores) representing the relative expression level of each AMPA subunit in the CR subpopulations (higher expression, red; lower expression, blue). (right) Expression of AMPA receptors displayed by UMI counts (y axis). Significant differential expression is indicated (***, Z-score>10).

The eight subclusters pointed to a yet undescribed heterogeneity within CR cells and to spatial and functional differences in the developmental pallium. We found *Cxcl12*, a chemokine secreted by the meninges and regulating the tangential migration of the CR cells^28^, to be also expressed by subtypes of CR cells (**Fig. 6b**). Notably, *in situ* hybridization data from E18 mice (Allen Brain Atlas) indicated that *Cxcl12+/Reln+* positive CR cells are located within the marginal zone (MZ), whereas *Cxcl12-/Reln+* are positioned outside the MZ, in the inner layers of the neocortex. Intriguingly, this points to a self-regulated migration capacity of the CR neurons of the marginal zone. The *bigSCale* analysis further unveiled potentially distinct differentiation stages of CR cells, marked by either *Sox11*/*Neurod2* or *Nnat*/*Igf2* (**Fig. 6a,c**). Likewise, we found a population of CR cells (CR8) expressing higher levels of mitochondrial genes, an indicator of apoptotic or disrupted cells (**Fig. 6a,c**). Considering that we did not find a similar cluster in the other pallial cell types, we excluded a technical artefact and suggest a cell subtype specific phenotype. Consistently, CR cells were shown to initiate cell death at postnatal stages^28^. Consequently, CR8 cells could represent an intriguing population of CR neurons committed to die already at the last stages of embryonic development (E18).

Lastly, neurotransmitter receptors are one of the most important features of CR cells. We specifically interrogated the expression of the 62 subunits of the 9 major receptor types. We found a number of subunits to be differentially expressed, pointing to CR subtypes with different membrane properties (**Supplementary Fig. 5**). The most striking variation was found for the Glu-R2 (*Gria2*), a pivotal subunit of AMPA channels strongly influencing receptor properties, assembly, trafficking and long-term synaptic plasticity (**Fig. 6d**).

## Discussion

Current scRNAseq analytic tools use simple or mixture probabilistic models which require predefined distributions to handle noise and sparsity. B*igSCale* bypasses this requirement by estimating a numerical model of noise. Furthermore, it determines the extent of the variation between cells without estimating actual gene expression value. These stratagems allowed us to build a highly optimized code, which can rapidly process large cell numbers whilst showing an improved sensitivity and specificity to detect differentially expressed genes, as shown for biological and simulated datasets. With the advent of microfluidic-based scRNAseq library preparation methods and the associated decrease in costs, experiments are now scalable to profile millions of cells simultaneously. Latest methods even provide single-cell transcriptomes without the physical separation of cells (through combinatorial indexing)^29^, paving the way to affordable big-scale projects and the comprehensive charting of tissue and organism compositions. With *bigSCale* we provide an analytical framework that addresses the computational challenges of future large datasets. While current tools are not applicable for experiments exceeding thousands of cells, DE analysis and clustering with *bigSCale* is practical for hundred thousand cells. Beyond that, its convolution module allows the analysis of millions of cells as shown here for the developing pallium.

With decreasing expenses for library preparation, sequencing costs become a limiting factor. Here we showed that despite being sequenced to low coverage (average 18,500 reads per cell), the analysis of more than a million cells is capable of identifying heterogeneity even in rare cell types. Indeed, the convolution into index cells and related improvements of expression profiles allowed us to draw a high-resolution atlas of the developing pallium, providing a rich resource of novel marker genes for subsequent studies. Further, the size of the dataset enabled us to describe a yet unprecedented heterogeneity in a rare, transient brain cell type (Cajal-Retzius neurons, 1% of total cells), producing new, founded hypotheses that can be used to enhance our mechanistic insights in brain development. Overall, these results illustrate the value of lowly sequenced large datasets. Nevertheless, for even sparser datasets, such as those obtained from the sequencing of nuclei^30^, the performance of *bigSCale* still needs to be evaluated.

Together, we present an analytical framework for scRNAseq analysis that provides a solution for challenges arising from future large-scale efforts to systematically and comprehensively chart cellular composition of complex organisms, including the human body^4^.

## Acknowledgements

We thank the cytometry unit of the CCiT (University of Barcelona) for successfully implementing single cell sorting. HH is a Miguel Servet (CP14/00229) researcher funded by the Spanish Institute of Health Carlos III (ISCIII). This work was further supported by the grants from FundaciÓn RamÓn Areces, the marathon “Todos Somos Raros, Todos Somos Únicos” (Proyecto 52). RC is recipient of a Marie SklodowskaCurie Actions fellowship (656359, H2020). Core funding is from the ISCIII and the Generalitat de Catalunya.

## Author’s contribution

HH and GI conceived the study. GI developed *bigSCale* and performed the statistical analysis. EM conceived and conducted the simulation analysis. AGA generated MARS-Seq sequencing libraries. GRE performed primary data processing. RC, IC, LAPJ contributed human NPC samples. MG and IG performed sequencing and provided computing infrastructure. HH and GI wrote the manuscript with support from EM. All authors read and approved the final manuscript.

## Methods

### Numerical probabilistic model

The probabilistic model is established as follows. First, cells are clustered in groups sharing similar expression profiles. We refer to this clustering as pre-clustering, as it is different from the final cell clustering achieved at the end of the pipeline. The purpose of the pre-clustering step is to group cells sharing highly similar transcriptomes, which are next treated as biological replicates to allow evaluation of the noise. Pre-clustering is achieved by *i)* normalizing the reads/UMIs to library size (*x*_*ij*_ *= c*_*ij*_*/LS for i=1,…,tot_genes*, where *x*_*ij*_ is the normalized expression for gene *i* in cell *j*, *c* the non-normalized expression and *LS=Sum(c*_*ij*_) the library size); *ii)* transforming the normalized expression levels in *log10(x+1)*; *iii)* normalizing the log-transformed values to the same interval for each gene. This step is required otherwise only highly expressed genes would drive the clustering; *iiii)* Clustering the cells using *Pearson*-correlation and hierarchical clustering with *Ward’s* linkage. *BigSCale* automatically attempts to find the deepest possible cut (on average *10-15%* of total tree height) in the tree to ensure that only highly similar cells are grouped together. At the same time, it avoids cuts which are too deep and would produce clusters which are too small for computing the numerical model. As a side note, the level at which the dendrogram is cut (and hence the number of pre-clusters) is not a key parameter of the pipeline, as it produces marginal effects on the final clustering or differential expression. Mainly, a lower number of pre-clusters will generate a numerical model which is less sensitive, meaning that the final p-values will be higher (less significant), but with negligible changes in the clustering and in the order of the differentially expressed genes. At this stage, we now treat the cells within each group as replicates, assuming their changes of expression to be solely due to noise and not to biological differences.

Secondly, all within-group pairwise comparisons between cells are enumerated in order to determine how rare/common (i.e. assigning a p-value) each combination of expression values is. Specifically, if a pre-cluster contains *n* cells, it produces *C(n,2)*=*n*(n-1)/2* combinations of cells. Each of this combinations contains *k* couples of expression values *(Xcell*_*1*_*, Xcell*_*2*_), where *k* is equal to the total number of genes and *Xcell*_*1*_*, Xcell*_*2*_ is the expression of a gene in the two compared cells. Each couple of expression values of each combination is summed into a 3D histogram that represents a numerical approximation of a cumulative distribution function (**Supplementary Fig. 6a,b**). The assigned p-values are related to the difference in gene expressions across all cells. For instance, if a gene has 0 UMIs in cell X and 2 UMIs in cell Y, its p-value would be larger than for a gene with 0 UMIs in cell X and 20 UMIs in cell Y, as such differences are rare.

The fitting takes into account the library size, meaning that it accounts for the higher dispersion of values of low-sized libraries. Specifically, when two cells of one pre-cluster are compared during the enumeration, they are normalized for the library size according to the formula *x*_*ij*_ *= c*_*ij*_*/Sum(c*_*ij*_*)*((LS*_*1*_*+LS*_*2*_*)/2)* for *i=1,…,tot_genes*, where *x*_*ij*_ is the normalized expression for gene *i* in cell j=*1 or j=2*, *c* the non-normalized expression and *LS*_*1*_*,LS*_*2*_ the library sizes of cells *j=1* and *j=2*).

Learning this numerical, probabilistic model from the data is possible because single-cell datasets contain hundreds to thousands of cells, which allows to enumerate up to hundreds of billions of couples and, hence, to gain a high precision in the estimated p-values. Ultimately, the model allows to assign a p-value to each gene, indicating the probability of a difference in the expression when comparing two cells.

### Differential expression (DE) model and hierarchical markers

The purpose of DE analysis is to assign p-values to genes that indicate the likelihood of an expression change between two groups of cells. To determine these p-values, each cell of one group is compared to each of the cells of the other group, resulting in a total of *n*_*1*_\**n*_*2*_ comparison, where *n* is the number of cells of each group. For each gene, the *n*_*1*_\**n*_*2*_ *log10* transformed p-values (derived from the probabilistic model and signed to represent up- or down-regulation) are summed into a total raw score. Genes up(down)-regulated in one group compared to the other will cumulate high (positive or negative) total raw scores. Here, the raw score is proportional to the likelihood of an expression change between the two groups.

The raw score is next adjusted *i)* for the total number of comparison, using a curve smoothing spline (**Supplementary Fig. 6c**). The rationale for this adjustment is to take into account that genes with sparser expression will produce smaller scores compared to genes expressed in high frequency; *ii)* for the within-group variability, which is estimated by running a DE analysis between randomly reshuffled cells in a way that cells of the same group are compared. Specifically, two null-groups are created by taking an equal proportion of cells from the two original groups. For example, in the case of two groups of 100 cells each, the null-groups will each be formed by mixed 50+50 cells randomly extracted from original group one and two, respectively. For comparison involving <2,000 total cells, 5 such permutation are performed. For comparison involving >2,000 total cells, the number of permutations is progressively scaled down with the increase of cell numbers. The reason is that large groups allow to fit the within group variability already with one or few permutations.

Aside from being a standalone tool, the DE script is also iteratively applied between clusters at the end of the clustering pipeline to isolate markers genes, i.e. genes expressed only in specific cells types (i.e. clusters). Upon completion of the clustering, a differential expression analysis is performed among all the pairs of clusters, resulting in (*N*^*2*^) comparisons, where *N=*number of clusters. Generally, the user can select the desired number of clusters, according to the desired detail of analysis. Nonetheless, *bigSCale* will calculate a hierarchical structure of the markers, which allows to recognize the main cell types even when setting a high N to inspect cell subtypes. In this way, the number of clusters N can be freely set to any level without the risk of losing phenotypic information.

As the last step, genes presenting significant changes of expression throughout the dataset are selected and organized in a hierarchical structure. Genes which are up-regulated in one population compared to each of the other populations are classified as markers specific to that population (Level 1 markers). Level 1 makers capture the phenotypes being unique and peculiar to populations of cells. Each Level 1 marker has a score, which corresponds to the highest (less significant) log10 transformed p-value out of the *N-1* comparison. In the next step, Level 2 markers are calculated. These markers are up-regulated in at most 2 populations of cells compared to each of the other populations. Essentially this means that Level 2 markers are genes expressed in two populations of cells amongst all populations. This computation iteratively continues up to Level *N-1* markers. Exemplarily, we assume four populations: radial glia, neuronal progenitors, dividing neuronal progenitors and differentiated neurons. Level 1 markers would represent genes expressed only in one of the populations, such as radial glia specific markers. Level 2 markers would be genes shared by two populations, such as the neuronal progenitors markers, which are expressed both in the neuronal progenitors and in the dividing neuronal progenitors. Lastly, Level 3 markers are shared by three populations, for example neuronal markers, which are expressed in the dividing and non-dividing progenitors and in the differentiated neurons.

To calculate new markers for CR cells we selected genes that were i) markers for CR cells, using the 1.3M convoluted dataset (1,291 genes with Z-score>6, **Supplementary Table 2**) and ii) uniformly expressed within subtypes of CR cells, using the deconvoluted CR dataset. Genes without significant changes of expression (max fold change <1.5) between subtypes (CR1-CR8) of CR cells were labeled as uniformly expressed in CR cells. A total of 501 genes including *Reln* satisfied both requirements. However, restricting the intersection to strong CR makers showing at least 8-fold increased expression in CR cells and Z-score>15 resulted in six high confidence makers: *Reln* (Z-score=35), *Cacna2d2* (Z-score=30), *Eya2* (Z-score=21), *Tex15* (Z-score=19), *Cpeb1* (Z-score=17), *Vmn2r1* (Z-score=15).

### Overview of the clustering

Once the probabilistic model has been fitted, it is possible to calculate distances between cells. Firstly, overdispersed genes, namely genes with high variation of expression throughout the dataset, are determined by means of a non-linear noise model learned from the data (**Supplementary Fig. 6d-f**). To further improve the features section, extremely skewed genes (**Supplementary Fig. 6g**) and isolated genes (not correlated with any others) are discarded. Furthermore, perfectly correlating genes are discarded as they belong to families with shared 3’-exons (such as *Pcdh* or *Uty*), for which most scRNAseq techniques (e.g. MARS-Seq^31,32^ or Chromium-based^3^ methods) cannot differentiate between transcripts. These families can otherwise generate artificial clusters, as it happens with other tools^15^.

Secondly, distances for all pairs of cells are calculated and the obtained distance-matrix is used to cluster the cells (hierarchical clustering, *Ward’s* linkage). The distance between two given cells is calculated as the sum of the log10-transformed p-values of overdispersed genes. Cells presenting many overdispersed DE genes will cumulate higher sums and eventually result very distant. Only genes with DE p-values<0.01 are retained in the sum to ensure that only significant changes determine the final distance. Prior to the calculation of the numerical model and distance matrix, batch correction can be applied to level out the batch-related variance in expression. Briefly, batch correction forces each gene to follow the same distribution in each batch, condition-wise (**Supplementary Fig. 6h**). In this way, the batch-effects are removed while preserving the original distributions of expression (**Supplementary Fig. 6h,i**).

### Convolution of large datasets

To convolute large dataset, *bigSCale* performs the following pipeline. 1) The numerical model of the dataset is calculated. 2) For each cell, its distances against a number n of other random cells are calculated. The number of random cells n is normally set to thousands. The higher n, the longer the computational time, but the lower the distortion introduced by the convolution. The final output of this step is a m*n matrix, where m is the number of cells in the original dataset and n is the number of random cells for which distances are calculated. 3) A pooling algorithm is applied to the *m*n* distance matrix to determine all groups of cells that will be summed into iCells. The rationale of the algorithm is that, for each cell, its closest neighbor among the *n* other random cells can be considered as an analogous phenotype. To increase the convolution factor, *k* closest neighbors, instead of 1, can be chosen. The pipeline pools the cells in order of similarity, starting with the closest ones, up to a maximum distance determined by percentile values. Initially, the algorithm starts with a stringent percentile value (*p=5%* of the total computed distances) and attempts pooling *k* closest neighbors for each cell. When there are no more cells with *k* closest neighbors within the maximum distance (*p=5%*), *k* is relaxed to *k-1*. This cycle continues until *k=1*, to which point the maximum distance allowed is increased to *p=10%*. These inner (*k*) and outer (*p*) cycle continue until *p=50%*. Cells ending up with no neighbors are considered outliers and discarded. While it is easy to locate neighbors for cells belonging to abundant (frequent) types, for rare cell types it becomes harder. Essentially, the two *k-p* cycles maximize the probability to find neighbors for every cell, both common and rare ones.

The ratio *n/k* is proportional to the quality of the convolution. In fact, a high *n/k* ratio implies that the k-closest neighbors chosen for each cell are selected from a much larger population of *n* random cells, which increases the chances to find the “real” neighbors, especially for rare cell types. Convolution of very large datasets can be split in multiple rounds to further reduce artifacts by using better *n/k* ratios, as done in the case of the 1.3M cells dataset (1.3 Million Brain Cells from E18 Mice, 10xGenomics). Specifically, we convoluted the dataset with a final factor k=75 in three rounds. In fact, calculating *n=4000* for 1.3M cells already requires approximately 12 hours of CPU-time, nonetheless yielding in a bad *n/k=4000/75=53,3* ratio, if convolution was in one round. Therefore, we proceeded with three rounds of convolution. The convolution factors used for each round where: (*n_1_=4000, k_1_=3*), (*n_2_=5000, k_2_=5*), (*n_3_=7000, k =5*) which all showed high, good *n/k* ratios (1333, 1000, 1400 respectively). The first round reduced the size to 456,274 iCells, the second round to 110,583 iCells and the third round to 26,185 iCells.

### Patient derived neural progenitor cells

Skin fibroblasts from two patients with Williams-Beuren (WB) and two with 7q11.23 microduplication (Dup7) syndrome were reprogrammed to induced pluripotent stem (iPS) cells by retroviral delivery of the pluripotency factors OCT4, SOX2, KLF4 and MYC, at the Centre of Regenerative Medicine in Barcelona (CMR[B]). Individual iPS cells were picked to generate single clone colonies that were expanded and fully characterized^33^. Briefly, genomic stability was confirmed by karyotype; integration and silencing were verified by PCR and quantitative RT-PCR; pluripotency was demonstrated by Alkaline Phosphatase staining and expression of pluripotency markers by immunocytochemistry. Finally, the capacity to differentiate to mesoderm, ectoderm and endoderm germ layers both *in vitro* and *in vivo* was verified by embryoid bodies and teratoma formation followed by immunostaining. All iPS cells were deposited in the Stem Cell Bank repository of the Instituto de Salud Carlos III ([SWB]FiPS1-R4F-5, [SWB]FiPS-4F-5-6, [DUP7]FiPS-4F-3-1, [DUP7]FiPS4-R4F-2). We generated neural progenitor cells from iPS cells following the Gibco protocol based on PSC Neural Induction Medium (NIM). Briefly, differentiation of iPS cell colonies was performed by seven days culture in NIM, followed by several passages of maturation in Complete Neural Expansion Medium (Neurobasal Medium, Advanced DMEM/F-12, and Neural Induction Supplement). Confirmation of expression of NPC markers was done by immunocytochemistry (Human Neural Stem Cell Immunocytochemistry Kit, Gibco). After 4-7 passages, NPC were detached with Accutase (Gibco) and resuspended. Single-cells were sorted in a BD Influx cell sorter to MARS-Seq plates (see below) for single-cell RNA sequencing (Flow Cytometry Core Facility, Univeritat Pompeu Fabra).

### Library preparation and sequencing

To construct single-cell libraries from polyA-tailed RNA, we applied massively parallel single-cell RNA sequencing (MARS-Seq)^31,32^. Briefly, single cells were FACS isolated into 384-well plates, containing lysis buffer (0.2% Triton (Sigma-Aldrich); RNase inhibitor (Invitrogen)) and reverse-transcription (RT) primers. The RT primers contained the single-cell barcodes and unique molecular identifiers (UMIs) for subsequent demultiplexing and correction for amplification biases, respectively. Single-cell lysates were denatured and immediately placed on ice. The RT reaction mix, containing SuperScript III reverse transcriptase (Invitrogen) was added to each sample. In the RT reaction, spike-in artificial transcripts (ERCC, Ambion) were included at a dilution of 1:16x10^6^ per cell. After RT, the cDNA was pooled using an automated pipeline (epMotion, Eppendorf). Unbound primers were eliminated by incubating the cDNA with exonuclease I (NEB). A second pooling was performed through cleanup with SPRI magnetic beads (Beckman Coulter). Subsequently, pooled cDNAs were converted into double-stranded DNA with the Second Strand Synthesis enzyme (NEB), followed by clean-up and linear amplification by T7 *in vitro* transcription overnight. Afterwards, the DNA template was removed by Turbo DNase I (Ambion) and the RNA was purified with SPRI beads. Amplified RNA was chemically fragmented with Zn2+ (Ambion), then purified with SPRI beads. The fragmented RNA was ligated with ligation primers containing a pool barcode and partial Illumina Read1 sequencing adapter using T4 RNA ligase I (NEB). Ligated products were reverse-transcribed using the Affinity Script RT enzyme (Agilent Technologies) and a primer complementary to the ligated adapter, partial Read1. The cDNA was purified with SPRI beads. Libraries were completed through a PCR step using the KAPA Hifi Hotstart ReadyMix (Kapa Biosystems) and a forward primer that contains Illumina P5-Read1 sequence and the reverse primer containing the P7-Read2 sequence. The final library was purified with SPRI beads to remove excess primers. Library concentration and molecular size were determined with High Sensitivity DNA Chip (Agilent Technologies). The libraries consisted of 192 single-cell pools. Multiplexed pools (2) were run in one Illumina HiSeq 2500 Rapid two lane flow cell following the manufacturer’s protocol. Primary data analysis was carried out with the standard Illumina pipeline. We produced 52 nt of transcript sequence reads.

### Data processing

The MARS-Seq technique takes advantage of two-level indexing that allows the multiplexed sequencing of 192 cells per pool and multiple pools per sequencing lane. Sequencing was carried out as paired-end reads; wherein the first read contains the transcript sequence and the second read the cell barcode and UMI. Quality check of the generated reads was performed with the FastQC quality control suite. Samples that reached the quality standards were then processed to deconvolute the reads to single-cell level by de-multiplexing according to the cell and pool barcodes. Reads were filtered to remove polyT sequences. Reads were mapped with the RNA pipeline of the GEMTools 1.7.0 suite^34^ using default parameters (6% of mismatches, minimum of 80% matched bases, and minimum quality threshold of 26) and the genome references for human (Gencode release 25, assembly GRCh38). Gene quantification was performed using UMI corrected transcript information to correct for amplification biases, collapsing read counts for reads mapping on a gene with the same UMI (allowing an edit distance up to 2 nt in UMI comparisons). Only unambiguously mapped reads were considered. The analysis of spike-in control RNA content allowed us to identify empty wells and barcodes with more than 15% of reads mapping to spike-in artificial transcripts were discarded. In addition, cells with less than 60% of reads mapping on the reference genome or more than 2x10^6^ total reads were discarded. Finally, low quality cells featuring either of the following were discarded: 1) low mapped reads 2) low library size 3) low library complexity (detected genes) 4) high mitochondrial content. Overall, 73 cells did not satisfy these quality requirements and were discarded.

### Simulated datasets

For data simulation we applied *Splatter*^12^ estimating parameters from NPC (*sim_NPC*) and a droplet-based experiment (2,520 random cells from: 1.3 Million Brain Cells from E18 Mice, 10x Genomics; *sim_10xG*). The datasets differed in the number of detected genes per cell, sparsity and heterogeneity. We recreated highly similar distributions of gene expression means and variances, cell library sizes and zeros counts as well as relationships of mean-variance and mean-zeros (**Fig. 2d** and **Supplementary Fig. 2a**). We preserved the number of cells and genes as in the original dataset and defined groups of different proportions across multiple sequencing pools. The dimensions of the gene x cell matrices were 41,020 x 1,847 and 27,998 x 2,520 in sim_NPC and sim_10xG, respectively. Each tool has been applied on the complete dataset at the model-building step, before defining groups of proportions 1:1 (1x), 1:2 (2x) and 1:10 (10x). The number of DE genes between groups ranged from 18% to 30% of the total number of DE genes (around 47% of total genes), being lowest at 10x and highest at 2x cases. While the composition of DE genes was similar in up-regulated and down-regulated genes, ratios of gene average means between groups could reach levels of expression magnitude up to twice as much as in sim_NPC. The datasets further differed in the proportion of outlier genes, which was around 1% in sim_10xG and ~2.5% in the sim_NPC.

ROC curves and pAUCs have been performed using the R package pROC^35^. In all comparisons, only genes tested by all methods were considered. Genes were ranked by nominal p-values, which we used to define a score as 1-p, indicating the outcome of the prediction (DE or non-DE) for each tool. Predictions and true gene labels were assessed at different thresholds of these scores to compute relative specificity and sensitivity coordinates for ROC curves.

### Data availability at GEO

The 1.3 million brain dataset is freely accessible from 10x Genomics:

<u>https://support.10xgenomics.com/single-cell-gene-expression/datasets</u>

The adult brain dataset is available at GEO (GSE60361).

The complete lists of hierarchical markers for the adult brain dataset^11^, the 10x Genomics dataset (1.3 Million Brain Cells from E18 Mice) and the *Reelin* subpopulations are available at GEO (GSE102934) in the following tables:

Table_S1_Linnarsson.xlsx (Markers of Zeisel/Linnarsson et al. dataset)

Table_S2_10xfull.xlsx (Markers of 10xGenomics)

Table_S3_10x_Reln.xlsx (Markers of 10xGenomics Reelin subtypes)

### Availability of the source code

All functions of *bigSCale* v1.0 are available at Github under the link: <u>https://github.com/iaconogi/bigSCale</u>

We are currently working to *bigSCale* 2.0, a user-friendly suite in which all parameters are automatically set and the analysis (DE and population clustering) can be performed in one-click.

